# Fine-mapping complex inversion breakpoints and investigating somatic pairing in the *Anopheles gambiae* species complex using proximity-ligation sequencing

**DOI:** 10.1101/662114

**Authors:** Russell B. Corbett-Detig, Iskander Said, Maria Calzetta, Max Genetti, Jakob McBroome, Nicholas W. Maurer, Vincenzo Petrarca, Alessandra della Torre, Nora J. Besansky

## Abstract

Chromosomal inversions are fundamental drivers of genome evolution. In the main afro-tropical malaria vector species, belonging to the *Anopheles gambiae* species complex, inversions play an important role in local adaptation and have a rich history of cytological study. Despite the importance and ubiquity of some chromosomal inversions across the species complex, inversion breakpoints are often challenging to map molecularly due to the presence of large repetitive regions. Here, we develop an approach that uses Hi-C sequencing data to molecularly fine-map the breakpoints of inversions 2Rbc and 2Rd in *A. coluzzii*. We found that inversion breakpoints occur in large repetitive regions, and strikingly among three inversions analyzed, two breakpoints appear to be reused in two separate inversions. Additionally, we use heterozygous individuals to quantitatively investigate somatic pairing disruption in the regions immediately surrounding inversion breakpoints, and we find that pairing disruption is undetectable beyond approximately 250 Kb from the inversion breakpoints.

## Introduction

Chromosomal inversions, reversals in the linear map order of chromosomes, are among the primary drivers of genome structure evolution across diverse species (Krimbas and Powell 1992; Hoffmann and Rieseberg 2008). Because they suppress recombination in heterozygous individuals, chromosomal inversions can maintain combinations of alleles that are more fit in similar contexts. Inversions are therefore theorized to be key contributors to local adaptation (Kirkpatrick and Barton 2006), speciation (Noor *et al.* 2001) and the maintenance of complex multigenic phenotypes (Lowry and Willis 2010; Joron *et al.* 2011). Owing to their myriad roles, uncovering the molecular and fitness consequences of inversions is a central goal for addressing numerous fundamental questions in evolutionary biology.

In the *Anopheles gambiae* species complex, inversions are known to play an important role in facilitating adaptation to a broad range of environments and to affect behavioural traits that may affect their efficiency as malaria vectors (Coluzzi *et al.* 1979; Rocca *et al.* 2009; Cheng *et al.* 2012; Ayala *et al.* 2014). Chromosome arm 2R, in particular, maintains a disproportionately large contingent of chromosomal inversions in the species’ genomes. Because this bias is evident in both common and rare inversions, it is thought to reflect a widespread mutational bias where inversions occur preferentially on this chromosome arm (Pombi *et al.* 2008). Furthermore, along 2R, specific cytological bands are strongly overrepresented for the presence or absence of inversion breakpoints, possibly consistent with mutational biases affecting the distribution of inversion breakpoints on short genomic scales as well (Coluzzi *et al.* 2002; Pombi *et al.* 2008). Uncovering the mutational processes that generate widespread chromosomal inversions in the *A. gambiae* species complex is key to understanding the ecological and evolutionary prospects for this group.

The precise identification and characterization of inversion breakpoints is a fundamental goal of evolutionary genomics. Breakpoint adjacent regions experience little or no recombination between arrangements and are particularly valuable for inferring the evolutionary histories of inversions (Wesley and Eanes 1994; Corbett-Detig and Hartl 2012), and provide ideal substrates for designing arrangement-specific PCR assays (*e.g. (Andolfatto et al. 1999; White et al. 2007; Lobo et al. 2010)*). Additionally, the genomic regions and specific structure of inversion breakpoints can yield key information about the molecular mechanisms underlying inversion formation, as well as the potential functional consequences of chromosomal inversions (Wesley and Eanes 1994; Puig *et al.* 2004). Nonetheless, precisely mapping inversion breakpoints at the molecular level is not always straightforward due to the presence of repetitive elements and large-scale duplications that are sometimes found in breakpoint adjacent regions of the genome.

Inversion breakpoint structures vary widely and impact prospects of successfully mapping the precise positions of inversion breakpoints. Whereas inversion breakpoints sometimes occur as simple “cut-and-paste” changes in unique sequences (*e.g.* those of *Drosophila melanogaster* and close relatives (Ranz *et al.* 2007; Corbett-Detig *et al.* 2012)), it is perhaps more common for breakpoints to occur in or to generate structurally complex regions often including repetitive elements (Cáceres *et al.* 1999; Lobo *et al.* 2010; Aguado *et al.* 2014). The former type of inversion breakpoint is relatively easily mapped using standard short-insert Illumina sequencing (*e.g.*, (Cridland and Thornton 2010; Corbett-Detig *et al.* 2012)). The latter can be particularly challenging to identify and often require the development of sophisticated molecular approaches (*e.g.*, (Aguado *et al.* 2014)).

In the *A. gambiae* species complex, some important inversion breakpoints have proven to be a persistent challenge for accurate breakpoint detection and assembly. In particular, (Lobo *et al.* 2010) used three Sanger assemblies (PEST, Pimperena, and Mali-NIH) together with directed BAC clone sequencing to accurately detect one of the breakpoints of 2Rb and one in 2Rbc, but were unable to identify the other breakpoints of either arrangement. The detected breakpoint contains a number of repetitive sequences, suggesting that this has been an important impediment to sequence-based detection of inversion breakpoints for these species. More recently, (Kingan *et al.* 2019) produced a *de novo* pacbio-based assembly of *A. coluzzii*. Despite reasonably high contiguity, we show here that their assembly fails to span important repetitive regions adjacent to known and our predicted inversion breakpoints. Thus, some of the inversion breakpoint adjacent regions in the *A. gambiae* species complex have been challenging to assemble using an array of genome sequencing technologies.

Proximity-ligation sequencing, or Hi-C, has recently emerged as a powerful method of detecting chromosome structure variation (Harewood *et al.* 2017; Himmelbach *et al.* 2018). Briefly, this technology enables one to sequence short reads from DNA molecules that existed close together in the chromatin of living cells, but not necessarily adjacent to each other in the primary chromosome sequence (Lieberman-Aiden *et al.* 2009). Importantly, Hi-C often produces read pairs that span large distances along a chromosome. Consequently, the complexity of breakpoint adjacent sequences has little impact on the ability to detect chromosomal inversions, but it is not always possible to resolve the breakpoint structures at the sequence level. Despite strong interest and several recent applications, there are few straightforward and automated approaches for basepair resolution characterization of structural variation breakpoints using Hi-C data.

One complication for the successful application of proximity-ligation sequencing for identifying inversion breakpoints is the presence of somatic chromosome pairing which is prevalent in Dipterans, including *A. gambiae* and *D. melanogaster* (Grell 1946). Somatic pairing occurs when homologous chromosomes are maintained in a physically attached state and consequently homologous alleles are in close physical proximity to each other in the interphase nucleus. This can be important for gene expression because enhancers on one paired chromosome may be able to initiate transcription of genes on the other, an effect known as transvection (Fukaya and Levine 2017). Inversions interfere with the somatic pairing process in heterozygotes, and may affect the pairing and allele proximity of breakpoint adjacent regions in heterokaryotypic individuals. Hi-C-based identification of inversion breakpoints is therefore likely to be impacted by somatic pairing and offer an opportunity to quantitatively investigate this phenomenon in a precise, high throughput format.

Here we apply Hi-C proximity-ligation sequencing to identify inversion breakpoints of 2Rc and 2Rd in *A. coluzzii.* We develop a simple approach for fine-mapping the positions of inversion breakpoints using Hi-C data and we use this to discover that all breakpoints in the inversions we study here occur in regions that contained repetitive elements before inversion formation. Because Hi-C assays sequence proximity within chromatin, this method also enabled accurate estimation of the potential extent of somatic pairing suppression due to structural heterozygosity. Strikingly, between just three inversions (c, d and b), there are only four unique breakpoint regions as two were reused twice. Our results suggest that unstable repetitive regions contribute disproportionately to inversion formation in the *A. gambiae* species complex.

## Results and Discussion

### Stocks and Sequencing Results

We obtained samples for homokaryotypic carriers of 2Rb, 2Rbc, and 2Rd arrangements (see Methods). For each colony or arrangement, we produced Hi-C libraries for pools of 5 whole adult mosquitos or 15-25 larvae following the library preparation protocol in (Lazar *et al.* 2018). We sequenced each library on a fraction of a Hiseq 4000 lane and obtained 12 million read pairs on average per sample. In each library, 23.5-36.8% of all read pairs mapped at distances of 1Kb or greater. Ultimately, we obtained relatively modest read depths (7.47X on average), but because of the long distances spanned between read pairs in Hi-C libraries, this corresponds to exceptionally high clone coverage (37,547X on average per site, Table S1).

### Fine-Mapping Approach

The nature of Hi-C data itself suggests a simple approach for mapping inversion breakpoint positions. We note that there are several methods for detection of structural variants from Hi-C (*e.g. (Harewood et al. 2017; Himmelbach et al. 2018)*). Nonetheless, to our knowledge, none of these have been validated for automated fine-mapping the specific locations of structural rearrangements. The primary reason is that by using a read binning strategy, previous automated approaches have placed a lower bound limit on breakpoint resolution.

We therefore developed, validated, and applied a simple alternative method of mapping inversion breakpoints that does not require read binning. Briefly, the key insight is that although Hi-C links often span long distances, the vast majority are still relatively short (Lieberman-Aiden *et al.* 2009). Therefore, if a sample has an inversion relative to the reference genome, this will artificially increase the apparent distances spanned by read pairs particularly in the regions surrounding inversion breakpoints. Importantly, even read pairs that map relatively distantly from the inversion breakpoints contain some information (albeit quite imprecise) about the locations of the breakpoints.

This observation suggests a simple approach for estimating inversion positions from the mapping positions of Hi-C short read data. Specifically, we seek to minimize the distance spanned by read pairs by transposing the mapping positions along a chromosome as defined by proposed breakpoint sites. We then optimize the joint position estimates using a Nelder-Mead direct search downhill simplex algorithm ((Nelder and Mead 1965), File S1).

### Validation

We validated this method using two previously mapped chromosomal inversions (2La and 2Rb). Both have been successfully characterized previously and both inversions are fixed within the *A. coluzzii* Mali-NIH colony that we used to identify the breakpoints of 2Rc (Sharakhov *et al.* 2006; Lobo *et al.* 2010). We therefore applied our method to these inversion breakpoints first, and we obtained strong concordance between the known inversion breakpoint position and our predicted mapping positions (Figure 1, Table S2). This suggests that this approach can accurately fine-map inversion breakpoints despite relatively modest sequencing read depths. (White *et al.* 2007; Lobo *et al.* 2010)

**Figure 1.**
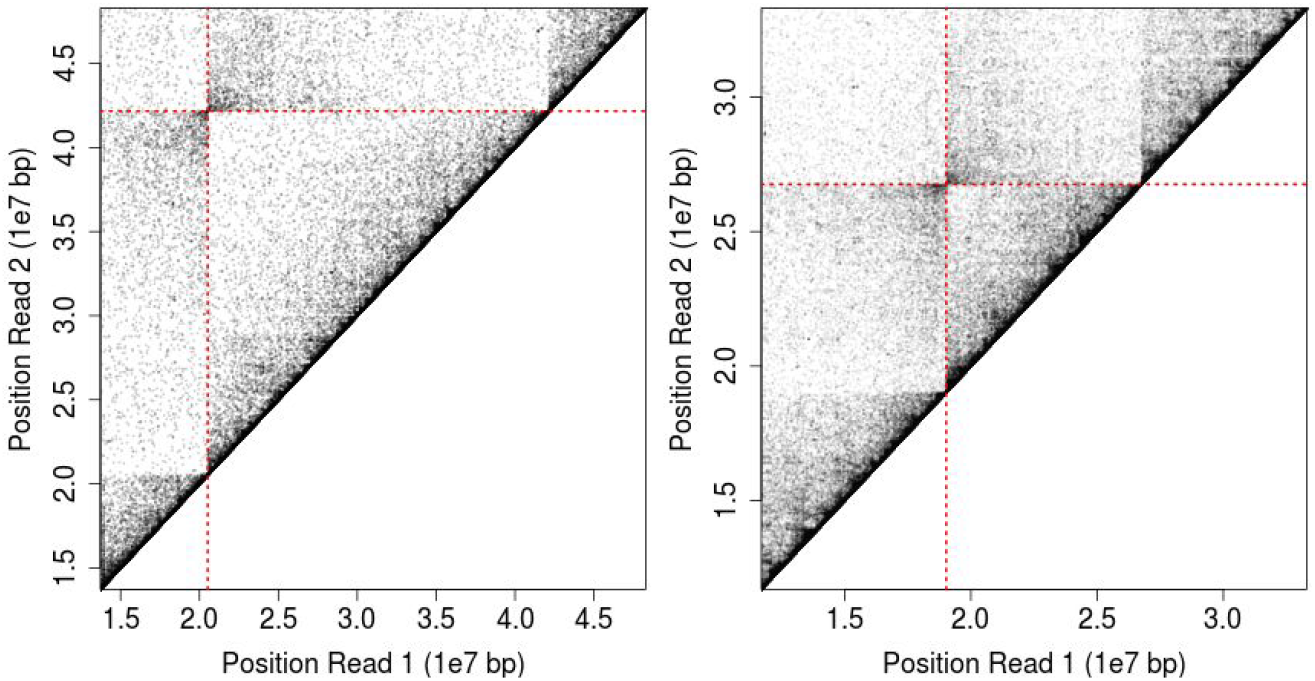
Validation of the fine-mapping Hi-C sequencing approach on *A. gambiae* inversions with known breakpoint positions. Mapping positions of Hi-C read pairs and predicted breakpoint positions (red lines) for 2La (left) and 2Rb (right) inversions.

### Robustness to lower read depths

We next sought to evaluate the robustness of our approach to a more modest sequencing effort. To do this, we subsampled the read pairs used to estimate inversion breakpoint positions focusing on inversion *2Rb* (Lobo *et al.* 2010). Despite light read coverage in many replicate subsampled sets (as low as ~0.2X mean read depth), we find that our method is able to consistently and accurately identify inversion breakpoint positions (Figure S1). This suggests that our approach can be applied even with relatively modest read depths and importantly that this method will be applicable even for extremely large genomes which could be cost prohibitive to sequence deeply using Hi-C or long-read technologies.

### Breakpoint Structures of *2Rc* and *2Rd*

We applied our approach to map the breakpoints of 2Rc and 2Rd in *A. coluzzii* and characterized the sequences surrounding each breakpoint. For both inversions, we found that all breakpoints localized to large annotated repeat clusters including both transposable elements and satellite repeat sequences in the standard arrangement AgamP4 reference assembly (www.vectorbase.org, (Giraldo-Calderón *et al.* 2015)). These regions are also often flanked by assembly gaps, suggesting that they have presented a persistent challenge for comprehensive genome sequencing and annotation. Because short read data cannot be accurately mapped within highly repetitive regions, we note that breakpoint estimates cannot be more accurately than localizing inversion breakpoints to within a specific repeat/gap cluster. Especially when a repeat cluster is relatively large, few or no reads will map uniquely within the repetitive region. Therefore breakpoint estimates will only be accurate to within the repetitive region identified but cannot precisely localized the breakpoint within the repeat cluster. Nonetheless, it is striking that each inversion breakpoint appears to be situated within large repetitive regions. In fact, the probability of selecting four regions at random along chromosome arm 2R with the same average rate of repetitive sequence annotation per basepair is small (P < 1e-4, Permutation Test), indicating that inversion breakpoint adjacent regions are strongly enriched for the presence of large blocks of repetitive sequences and assembly gaps.

It is also noteworthy that two repetitive regions appear to be reused between just these three inversions—an extremely improbable event by chance (P < 1e-4, Permutation Test; see Methods). Specifically, we find reused breakpoints between the proximal breakpoint of 2Rb and the distal breakpoint of 2Rc, and between the proximal breakpoint of 2Rc and the distal breakpoint of 2Rd (Figure 2, Table S2). Previous work supports the shared breakpoint positions of 2Rc and 2Rd which are cytologically indistinguishable, but not those of 2Rb and 2Rc, because of the presence of a thin band between the two breakpoints (Figure S2), suggesting that a relatively large genomic segment lies between them (Coluzzi *et al.* 2002). Nevertheless, it needs to be highlighted that, although the exclusion of the thin band between 2Rb and 2Rc inversions was considered the most probable interpretation during the creation of the polytene chromosome map, cytological interpretation leaves some elements of obvious uncertainty. Alternatively, given the demonstrated accuracy of our mapping approach, one possible explanation is that the breakpoints are close in reference coordinates but that the reference has a large gap, possibly due to the presence of large intervening collapsed repeat sequences.

**Figure 2.**
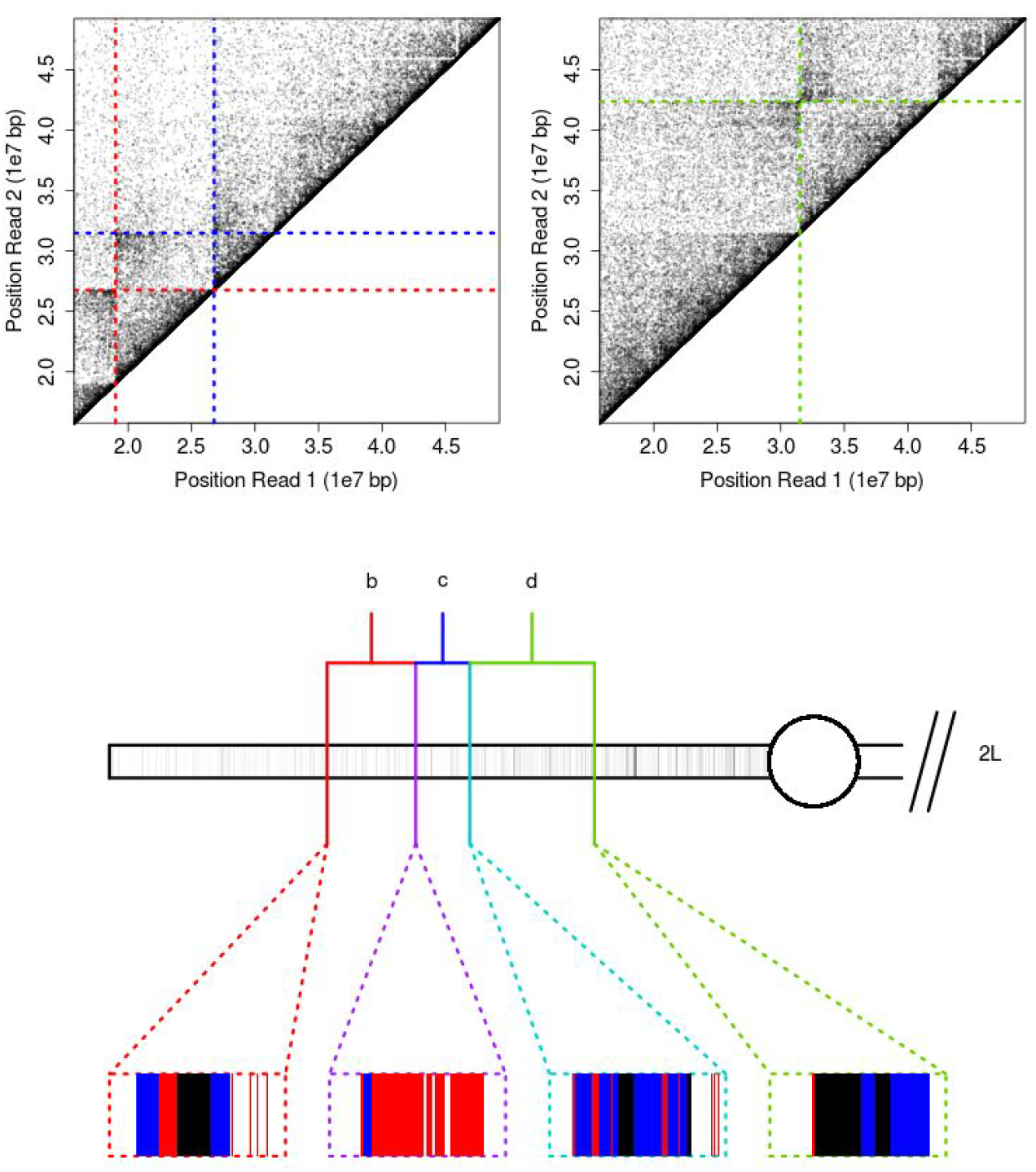
Breakpoint positions of inversions 2Rb, 2Rc and 2Rd in *A. coluzzii* and schematic of breakpoint adjacent sequences. The breakpoint mapping positions for individuals with arrangement 2Rbc (top left) with predicted breakpoints indicated for 2Rb (red) and 2Rc (blue). Breakpoint mapping positions for individuals with arrangement 2Rd (top right), with predicted breakpoint positions shown in green. A schematic of chromosome arm 2R (bottom) with positions of inversions indicated and a breakpoint structure schematic of 40 Kb surrounding each breakpoint. This schematics include satellite repeat sequences (blue), assembly gaps (black), and other repeats (red). Repeat annotations are from vectorbase.org and exclude all repeats of less than 100 bp in length.

Breakpoint reuse at the molecular level may have two causes. First, it is possible that specific genomic regions have higher relative fitness with inversion breakpoints. This could be either because these regions are more fit as a consequence of structural rearrangements (*e.g.*, if they constitute pairing sensitive sites, (Corbett-Detig 2016)), or because these regions suffer a lower fitness cost due to the presence of large structural rearrangements. Alternatively, these regions may harbor genes that recurrently contribute to ecological differentiation among *Anopheles* species. In support of this hypothesis, Coluzzi (Coluzzi *et al.* 2002) previously observed that this region on 2R is frequently rearranged during *Anopheles* evolution and suggested that this region contributes to adaptive differentiation associated with oviposition site—a fundamental ecological characteristic of these species.

Second, if these regions are simply more prone to breakage due to higher intrinsic fragility, breakpoint reuse could be expected as a consequence of neutral processes (Krimbas and Powell 1992; Caceres *et al.* 1997). The extensive distribution of satellite sequences in breakpoint adjacent regions is consistent with the second explanation and suggests that breakpoint reuse in at the molecular level in the *A. coluzzii* species complex is a consequence of higher rates of breakage in these specific regions. Nonetheless, the accumulation of transposable elements near many breakpoints also suggests that structural heterozygosity was readily tolerated in these regions prior to inversion formation and therefore that fitness costs of rearrangements are minimal.

### Comparison to a Long-Read Based Assembly

Recently, (Kingan *et al.* 2019) produced a *de novo* genome assembly for *A. coluzzi* using high coverage pacbio long read sequence data. This colony bears the same arrangement as the AgamP4 reference genome. To determine if their approach could assemble across these large-scale repeats and thereby reveal the molecular organization of the breakpoint associated regions, we aligned the genome to the AgamP4 genome assembly and extracted the contigs that aligned adjacent to each large-scale repeat cluster. For all three putative breakpoints, we found large contigs (all greater than 500 Kb) that aligned collinear to the breakpoint adjacent regions in the *A. gambiae* genome assembly. However, we did not identify a scaffold that spanned any of the predicted breakpoints (Table S3), indicating that these genomic regions remain a persistent challenge for even the most advanced long read sequence-based assembly methods. In fact, a single contig spans the length of the genomic segment that contains 2Rc and terminates on each end at the repeat clusters surrounding our predicted inversion breakpoints. Nonetheless, this does reinforce a key advantage of Hi-C-based inversion breakpoint detection. Specifically, chromatin conformation capture can span very large genomic distances thereby mitigating the impacts of large repetitive regions which may be challenging or impossible to completely sequence.

### Repeat Content Is Unlikely to Explain Mutational Biases Among Chromosome Arms

Although repetitive elements may play an important role in the generation of inversion breakpoints for the three inversions investigated here, the distribution of repeats across the *Anopheles* genome is unlikely to explain the abundance of breakpoints on chromosome arm 2R (Pombi *et al.* 2008). Across each autosomal chromosome arm in the most recent assembly of *A. gambiae*, AgamP4, chromosome arm-2R has a relatively low rate of annotated repetitive elements (5.3% of sites on 2R are annotated as repeats longer than 1 Kb, versus 5.3-6.3% across 2L, 3L, and 3R). Similarly, 2R does not contain an excess of satellite repeat elements specifically (0.09% of sites on 2R are annotated as satellites, versus 0.09-1.8% on 2L, 3L, and 3R). Alternative mechanisms beyond a simple abundance of repetitive sequence is therefore more likely to explain the proliferation of rearrangements on 2R specifically. Nonetheless, it is possible that arm 2R contains a disproportionately large amount of unassembled repeats in the *A. gambiae* genome, thereby obscuring this effect.

### Impact of Somatic Pairing on Breakpoint Identification

Whereas sister chromosomes in mammalian genomes maintain independent chromosome domains in somatic tissues, dipteran sister chromosomes are paired along their lengths in the vast majority of somatic cells (Metz 1916). Heterokaryotypy is therefore expected to impact our prospects for successfully mapping inversion breakpoints. To investigate this phenomenon, we produced and sequenced an additional library from 2Rd/2Rd+ heterokaryotypic individuals. Whereas the homozygote library reveals a strong enrichment for Hi-C links in the lower left and upper right quadrants (Figure 3), the heterokaryotype library is much less strongly delineated. When we attempt to bioinformatically map the breakpoints as described above, our method fails, presumably due to the challenges associated with somatic pairing.

**Figure 3.**
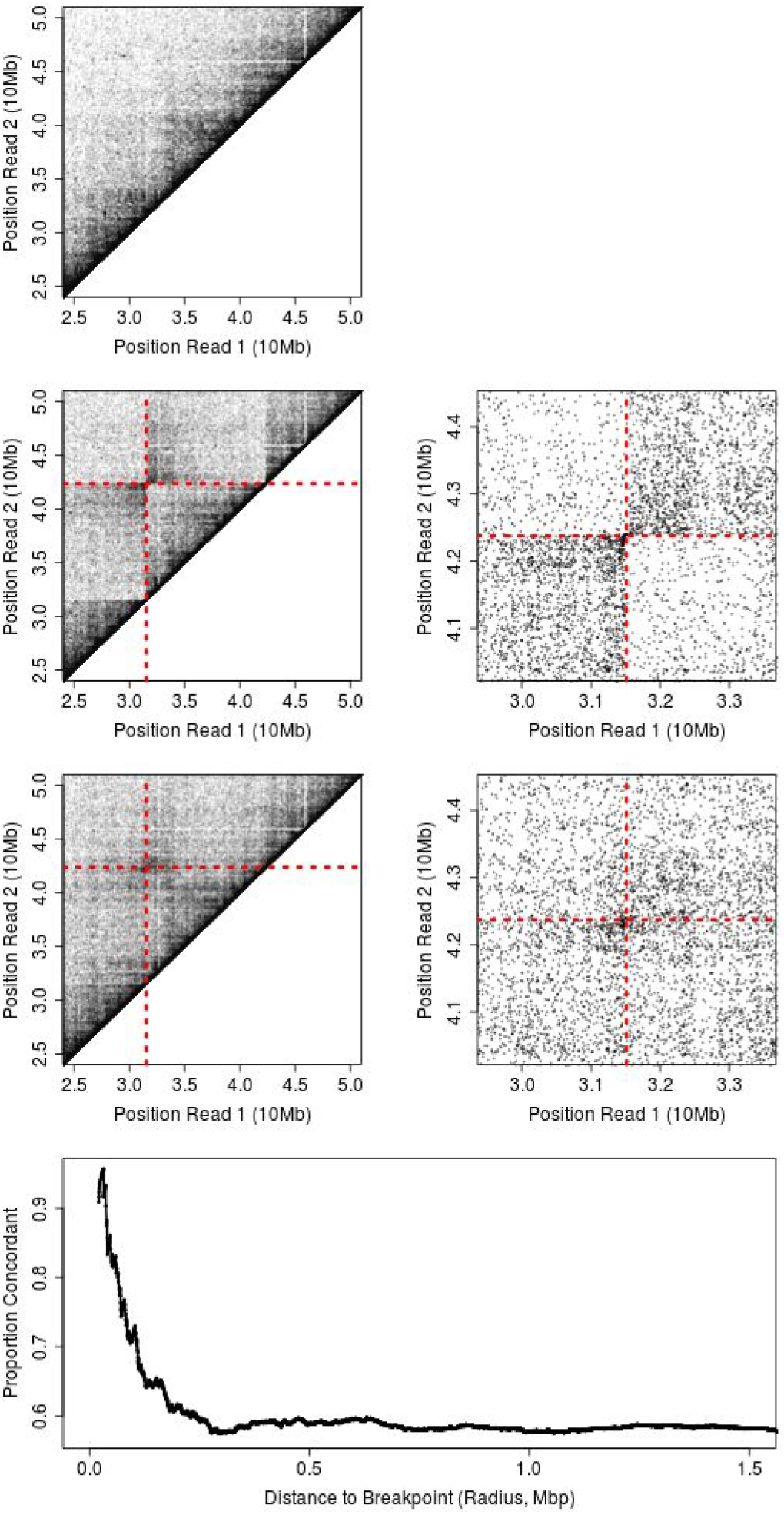
Impact of somatic pairing for 2Rd/2Rd+ heterokaryotypic *A. coluzzii*. For reference, the contact map of a standard arrangement, homokaryotypic individual (top). Inversion breakpoint predictions and contact map for homokaryotypic individuals (2nd row, left), and the 4 Mb window surrounding the breakpoint region in homokaryotypic individuals (2nd row, right). Inversion breakpoint predictions and contact map for heterokaryotypic individuals (3rd row, left), and the 4 Mb window surrounding the breakpoint region in heterokaryotypic individuals (3rd row, right). Finally, the proportion of read pairs that are concordant with the inversion structure (i.e. map into the lower left or upper right quadrants), with increasing distance from the inversion breakpoint, where distance is measured as a straight line between the breakpoint and coordinates defined by each read pair.

To attempt to map breakpoint positions in heterokaryotypes, we modified our mapping approach to accept only read pairs for which the first is within 5 Mb of the distal breakpoint and the second is within 5 Mb of the proximal breakpoint. In rerunning our mapping approach, the distal breakpoint estimated position is predicted at position 31,495,608, which is remarkably close to our estimate from homokaryotypic individuals and within the same repetitive sequence block. However, the proximal breakpoint is predicted at position 42,550,800 which is approximately 175 Kb from the breakpoint we predicted from homokaryotypic individuals. This difference may reflect the challenges of the real chromatin domains, which alter the frequencies of links and suggests that whenever feasible, homokaryotypes should be used for mapping breakpoint positions when working with dipterans or other species that experience somatic pairing.

### Breakpoint Heterozygosity and Somatic Pairing

Despite the diffuse signal of association between sister chromosomes, there is still weak enrichment for the lower left and upper right quadrants (28% and 30% of read pairs respectively within a 4 Mb square centered on the breakpoint) in the 2Rd/2Rd+ heterokaryotype Hi-C mapping data (Figure 3). Read pairs mapping in these quadrants correspond to those that are physically proximal along the inverted chromosome. This suggests that maternal and paternal chromosomes are almost equally likely to contact each other as to contact themselves even in the regions relatively near to breakpoints and that inversion breakpoints present little barrier to somatic pairing despite different chromosome structures on broad scales (similar to inversions in *D. melanogaster (Golic and Golic 1996)*).

To evaluate the impact on somatic pairing more quantitatively, we computed enrichment for contacts in the lower left and upper right quadrant with increasing radius outward from the point defined by the 2Rd inversion breakpoints. Our results indicate that read pairs that are extremely close to the inversion breakpoints are much more likely to contact the same chromosome, but that this effect decays quickly to background levels within approximately 250 Kb (Figure 3). Our data therefore provide a quantitative estimate of the potential scale of impacts of structural heterozygosity on somatic pairing and therefore on interchromosomal effects on gene expression.

Transvection is a phenomenon where sister chromosomes affect expression of their homologs and plays a potentially fundamental role in genome evolution (Geyer *et al.* 1990; Duncan 2002). Transvecting enhancers are able to contact and initiate transcription of homologous target genes in trans on paired chromosomes, allowing rescue of a haplotype with a nonfunctional enhancer (Mellert and Truman 2012). The process requires somatically paired homologs to share transcriptional machinery in the form of chromatin-affecting proteins and transcription factors. Our data indicate that pairing is suppressed in the immediate vicinity of the breakpoints in heterokaryotypes, which would also suppress transvection effects (Golic and Golic 1996). Transvection would be unaffected in homokaryotypes since their pairing appears largely normal (Figure 3). However, since the majority of genomic elements immediately surrounding inversion breakpoints are repetitive sequences, it is unlikely that heterokaryotypy strongly impacts gene expression by preventing potential transvection effects for 2Rd heterozygotes and for other inversions across the species’ range more broadly.

## Conclusion

Accurate inversion breakpoint detection is central for evolutionary genomic inference and for developing molecular karyotyping diagnostics. Here we have shown that Hi-C sequencing is a cost-effective means of accurately fine-mapping inversion breakpoints in members of the *Anopheles* species complex. Our results demonstrate that conventional binning approaches for analyzing Hi-C contact maps are not a prerequisite, and limitations imposed by these methods can therefore be avoided even for samples with very modest sequencing depths. Importantly, Hi-C has virtually unlimited range despite extensive repetitive sequences flanking the inversion breakpoints of interest in the *A. gambiae* species complex. Breakpoint identification reliant on Hi-C data and related approaches will therefore enable structural variation discovery across the *A. gambiae* species complex as well as across life more generally.

Recent work in *A. gambiae* and *Drosophila* species has found chromosomal inversion structure to have little effect on gene expression patterns (Huang *et al.* 2015; Fuller *et al.* 2016; Lavington and Kern 2017; Said *et al.* 2018; Cheng *et al.* 2018). Our observation that somatic pairing is disrupted only partially and only in the relatively small regions immediately surrounding inversion breakpoints is consistent with these observations and suggests that dipteran chromatin structures are particularly resilient to changes imposed by genome structure polymorphism. Our results in conjunction with previous molecular work (Golic and Golic 1996) contribute to a growing understanding of extremely abundant chromosomal structural heterozygosity within clades of dipteran insects.

## Methods

### Anopheles gambiae s.l. Colonies

We obtained adult or larval mosquitoes of the Pimperena, Mali-NIH, and Ndokayo colonies from BEI Resources (*Anopheles* program; https://www.beiresources.org/AnophelesProgram/Anopheles/WildStocks.aspx) *A. gambiae* Pimperena (2Rb), *A. coluzzii* Mali-NIH (2Rbc), and *A. coluzzii* Ndokayo (2R+^bc^, i.e. standard orientation for 2Rbc arrangement). In addition, carcasses of homokaryotypic and heterokaryotypic 2Rd carriers were selected by cytological analysis of ovarian polytene chromosomes (Torre and della Torre 1997) of half-gravid females from 2Rd-polymorphic *A. coluzzii* Banfora M colony (Liverpool School of Tropical Medicine and Hygiene, LSTMH, UK), established from samples collected in 2014 from Banfora District, Burkina Faso, by LSTMH with support from the Centre National de Recherche et de Formation sur le Paludisme (CNRFP, Burkina Faso). Samples were kept at −80°C until library preparation.

### Hi-C Library Preparation and Sequencing

To extract nuclei, we placed five adult mosquitoes into a dounce homogenizer and we used 5-10 strokes of the pestle to homogenize the contents. We then filtered the homogenate through a 40um screen and fixed nuclei with 1% formaldehyde. We produce Hi-C libraries as described in (Lazar *et al.* 2018), and we sequenced each library on a portion of a Hiseq 4000 lane.

### Read Mapping and Filtering

After chromatin capture on steptavidin beads, we sheared DNA with a restriction enzyme, end repaired, and blunt-end ligated the end repaired fragments. This causes junctions between fragments to be demarcated with two intact copies of the enzyme’s recognition sequence. In this case “GATC”. We therefore searched all reads for the characteristic “GATCGATC” sequence and replaced that sequence plus any remaining sequence on the read with a single GATC, which must have been present in the sequence.

We then mapped trimmed short read data to the AgamP4 *A. gambiae* reference genome using BWA v0.7.17 using the mem alignment function. We filtered all reads with a mapping quality of less than 30 and we removed all reads whose pairs did not successfully map to the reference genome.

### Breakpoint Position Estimation

We estimated 2Rc and 2Rd inversion breakpoint positions as a two-parameter optimization task. Specifically, we seek to minimize the total distance spanned by all read pairs surrounding inversion breakpoints by inputting possible breakpoints positions, “reversing” the inverted region, and recomputing the distance spanned by all read pairs. We implemented this procedure in python (FIle S1) and used the scipy optimize() function to implement a Nelder-Mead (Nelder and Mead 1965) two parameter optimization procedure. We evaluated the accuracy of our approach by comparing our estimated breakpoint positions for 2La and 2Rb, which have been identified previously (Sharakhov *et al.* 2006; Lobo *et al.* 2010), and we evaluated the robustness by randomly subsampling read pairs and re-estimating inversion breakpoints at increasingly small read depths for 2Rb.

### Permutation Tests

We tested for an enrichment for large blocks of repetitive sequences adjacent to 2Rb, 2Rc and 2Rd inversion breakpoints using a permutation test framework. Specifically, we randomly drew positions for the four breakpoints from all positions on 2R. We then computed the proportion of sites annotated as repetitive or assembly gaps within surrounding 40 Kb windows, and we asked if the mean repetitive/gap sequence content equalled or exceeded the amount we obtained from the true breakpoint positions. We then recorded the proportion of replicates that satisfied these criteria.

We also used a permutation test to ask if the breakpoint co-localization among separate inversions could be expected by chance. Here we assume that all inversion breakpoint are sampled independently from the chromosome arm. To accommodate our uncertainty with the exact breakpoint positions within large repetitive blocks of sequence, we recorded two breakpoints as co-localized when they coincide to within the same block of repetitive sequence or within 5 Kb in coordinate space if we did not draw a position within an annotated repetitive region. We then asked if each replicate permutation produced two or more colocalized breakpoints and recorded the proportion of such tests.

We performed each permutation procedure 10,000 times.

### Data Availability

All sequence data produced in this work will be available from the sequence read archive under project accession number PRJAXXX.

## Acknowledgements

We thank E. Dotson, Principal Investigator for MR4 Vector Activity, for kindly supplying Mali-NIH, Pimperena, and Nkdokayo mosquitoes, and H. Ranson for sharing the Banfora M colony.

Support for this work came from the National Institutes of Health (R01 AI125360 awarded to NJB and R35 GM128932 to RBC-D) and from a Alfred P. Sloan Fellowship to RBC-D. During this work JM and MG were supported by NIH training grant T32 HG008345-01.

---

**Figure S1.**
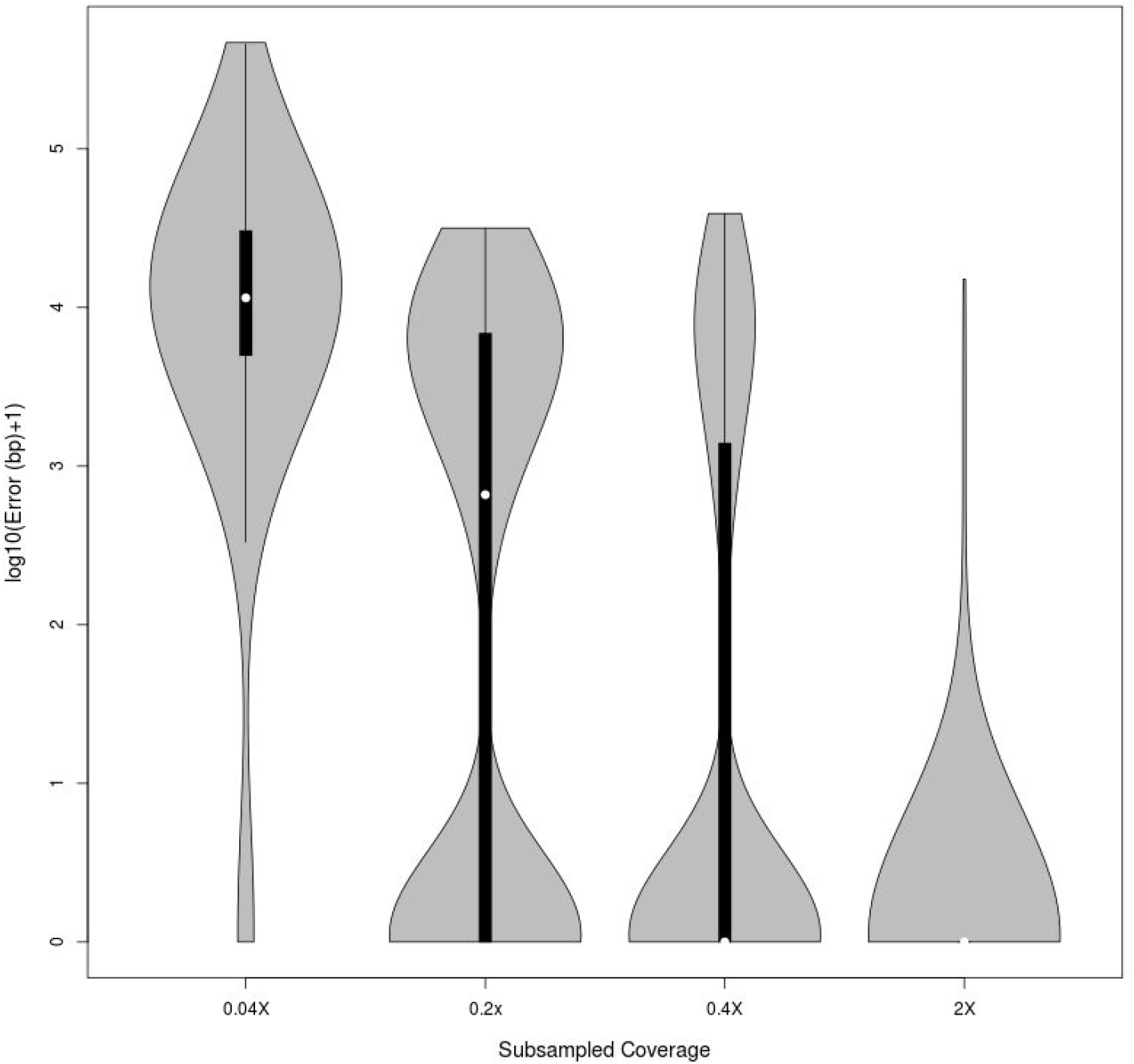
Error in breakpoint position estimates at subsampled lower read depths. Violin plots of the distribution of error in breakpoint estimated positions for inversion 2Rb for 1,000 replicate subsampled sets at depths 0.04X, 0.2X, 0.4X and 2X.

**Figure S2.**
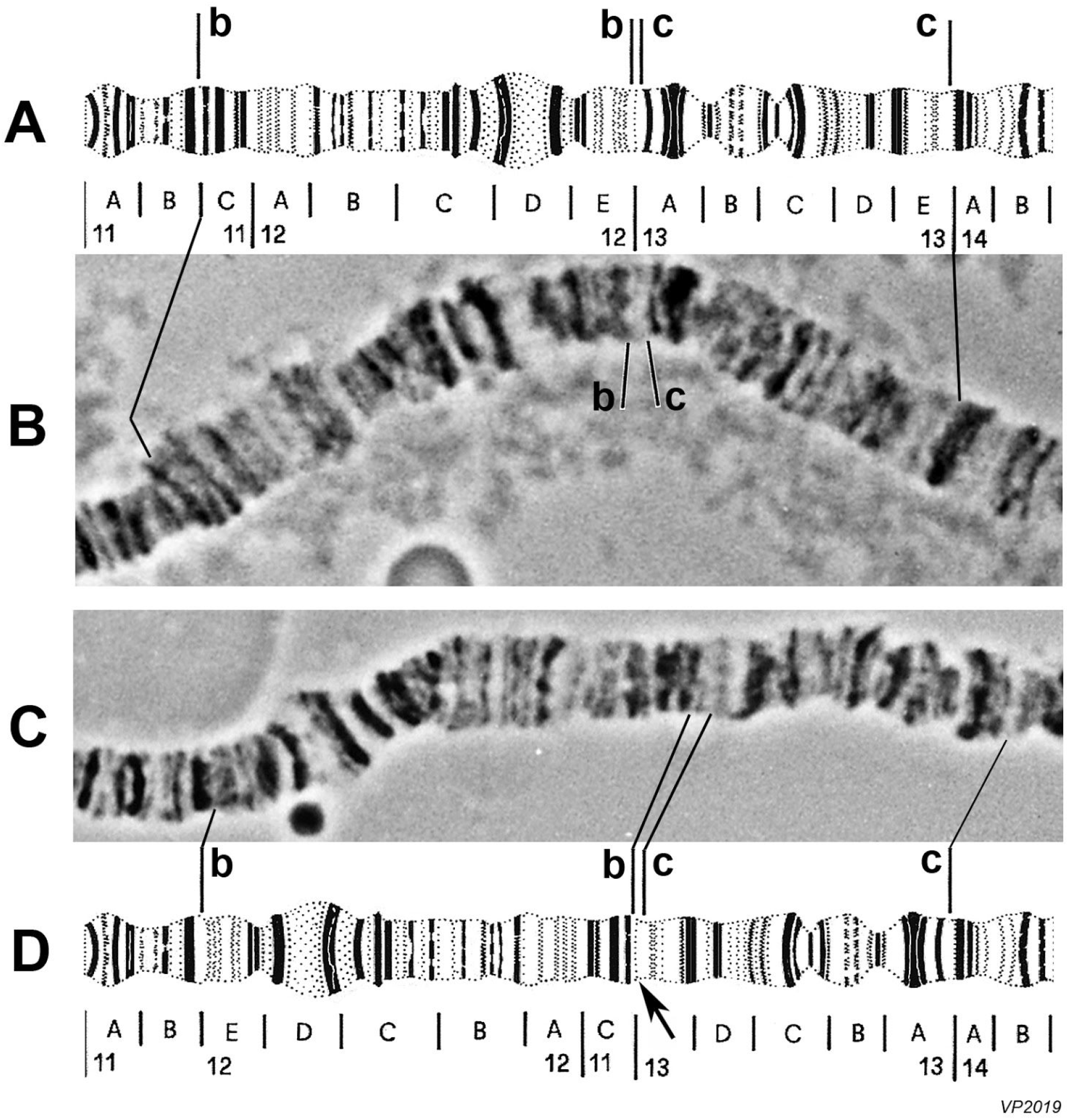
Polytene chromosomal arm 2R of *Anopheles gambiae*, subdivisions and divisions 11 to 14 (partim). A: diagrammatic uninverted (standard) arrangements of inversions 2Rb and 2Rc; B: polytene 2R +^b^/+^b^ -- +^c^/+^c^ standard homokaryotype; C: 2Rb and 2Rc inverted homokaryotypes (2Rbc/bc); D: diagrammatic 2Rb and 2Rc homokaryotypes (2Rbc/bc). Thin lines indicate useful map landmarks. Arrow point to the thin band (first of subdivision 13A) believed to be excluded from both inversions 2Rb and 2Rc. *Anopheles gambiae* map diagrammatic representations modified from Figure 1 and poster in Coluzzi et al, 2002. Picture B: *An.gambiae* from South Africa; picture C: *An.gambiae* from Jirima (Kano State), North Nigeria.

**Table S1.** Library properties for Hi-C data collected in this work.

| Colony | Arrangement | Read Pairs | Read Coverage | Proportion Mapping > 1Kb | Mean Clone Coverage |
| --- | --- | --- | --- | --- | --- |
| Ndokayo | +/+ | 8006299 | 4.31X | 0.368 | 19378.9 |
| Pimperena | 2Rb/2Rb | 7463895 | 4.01X | 0.334 | 16159.9 |
| Mali-NIH | 2La/2La;<br>2Rbc/2Rbc | 1945753 | 1.12X | 0.235 | 4511.0 |
| Banfora M | 2Rd/2Rd | 19098259 | 12.25X | 0.310 | 61008.6 |
| Banfora M | 2Rd/2R+d | 24965090 | 15.68X | 0.307 | 86678.9 |

**Table S2.**
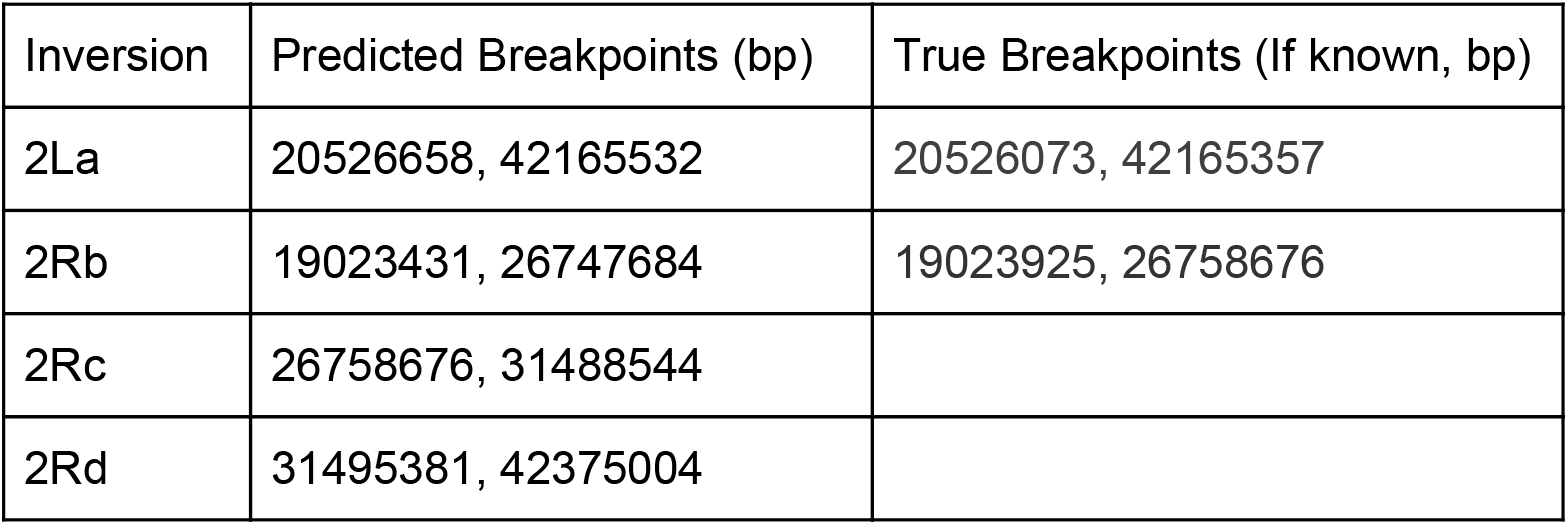
Predicted mapping positions of inversions studied in this work.

**Table S3.**
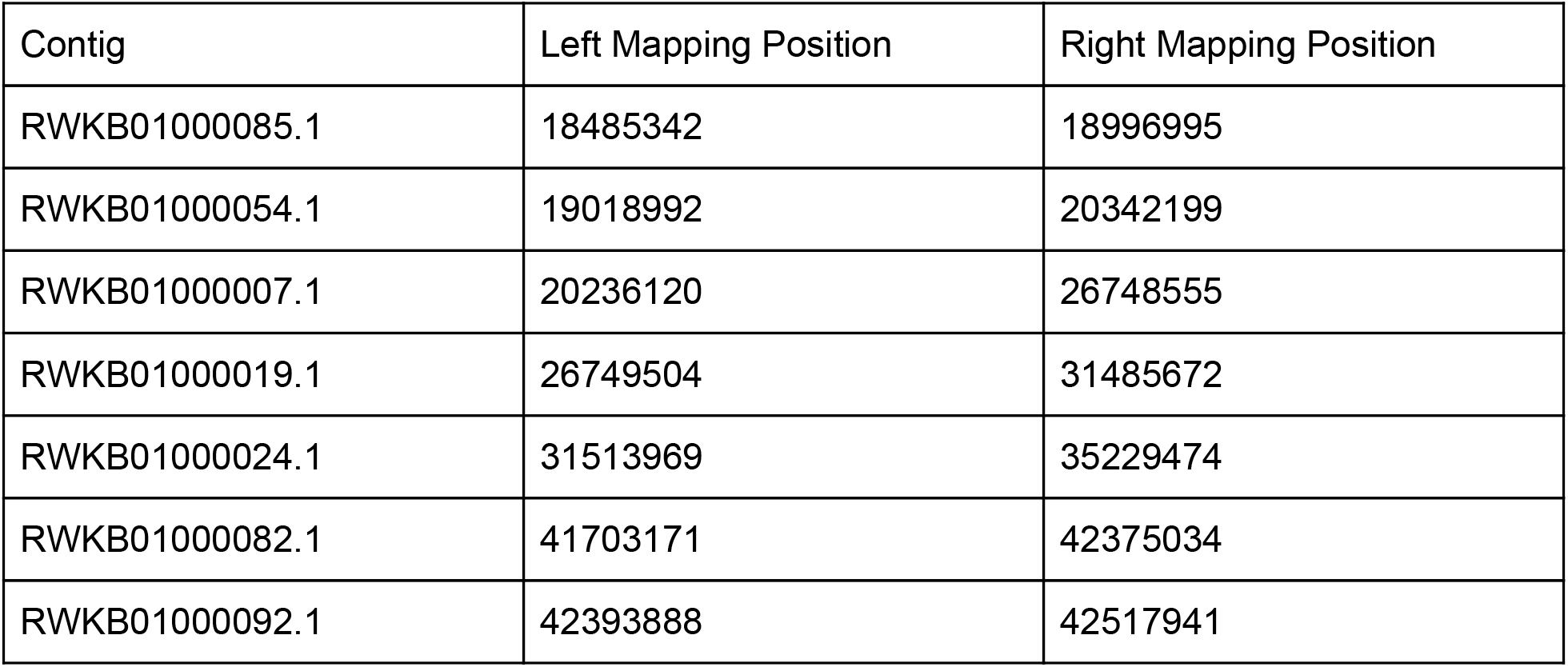
Breakpoint adjacent contigs obtained from a recent *A. coluzzii* genome assembly.

| Contig | Left Mapping Position | Right Mapping Position |
| --- | --- | --- |
| RWKB01000085.1 | 18485342 | 18996995 |
| RWKB01000054.1 | 19018992 | 20342199 |
| RWKB01000007.1 | 20236120 | 26748555 |
| RWKB01000019.1 | 26749504 | 31485672 |
| RWKB01000024.1 | 31513969 | 35229474 |
| RWKB01000082.1 | 41703171 | 42375034 |
| RWKB01000092.1 | 42393888 | 42517941 |

**File S1.**
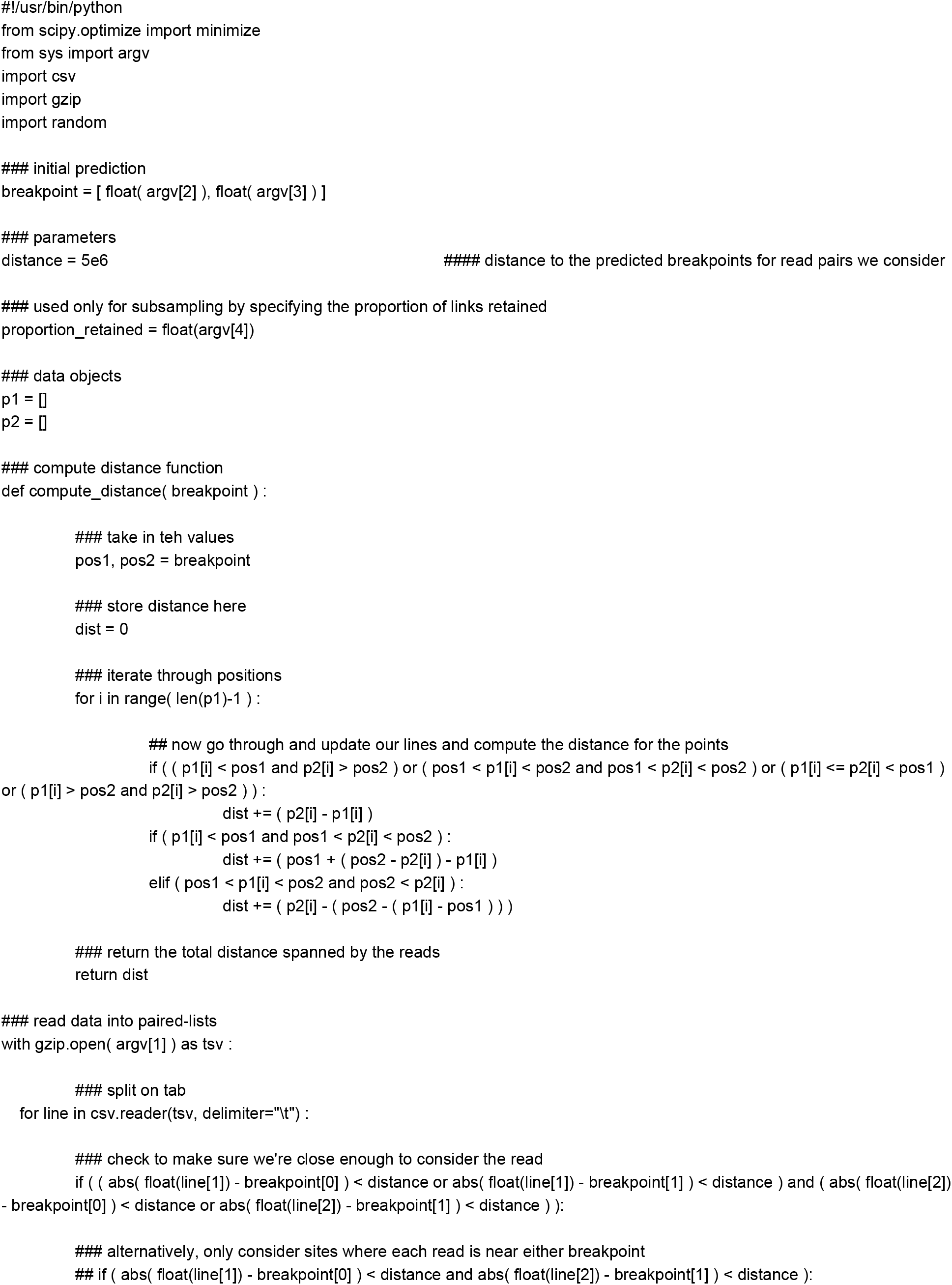
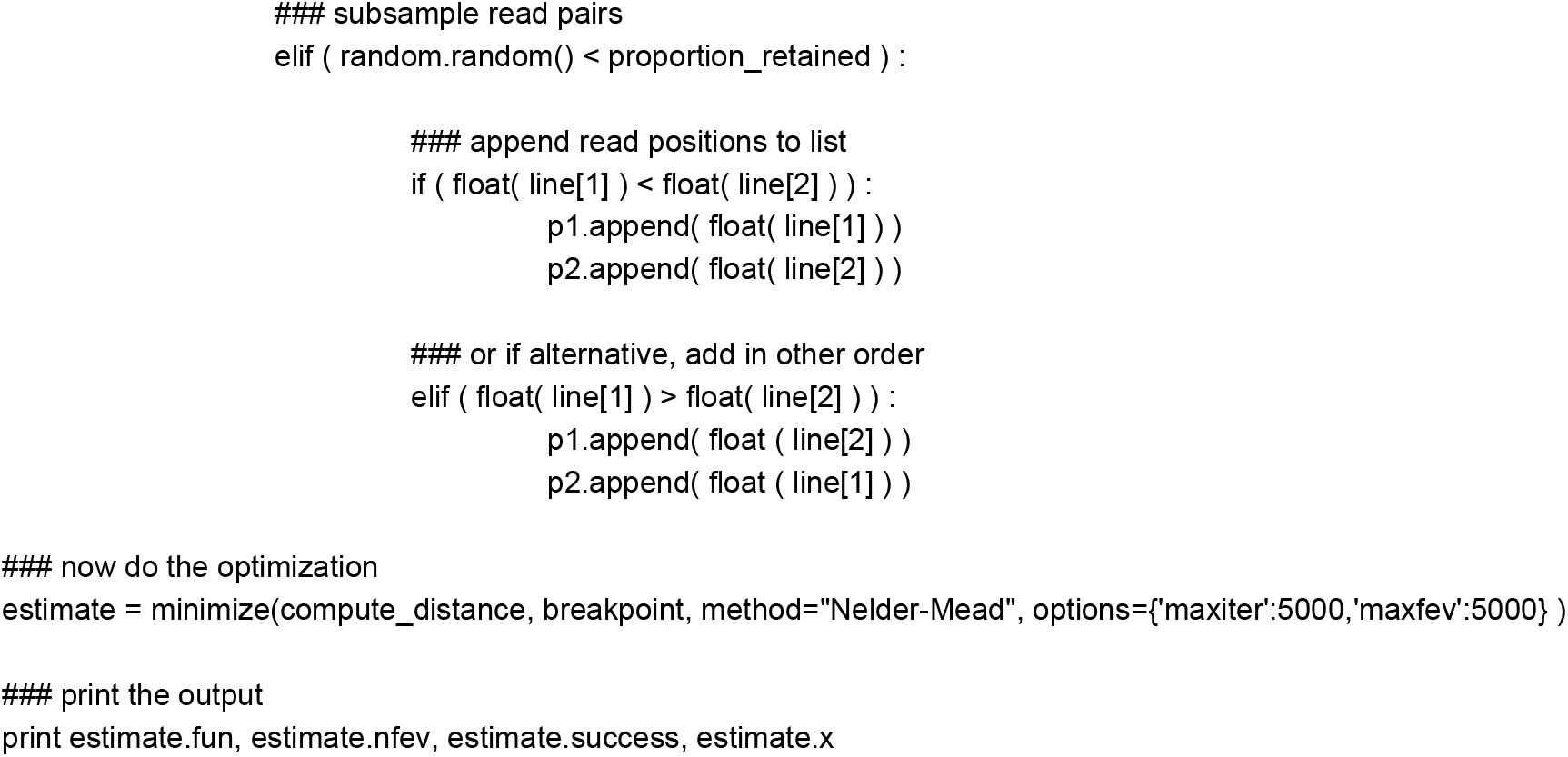
Script used to optimize breakpoint position estimates.

